# Structure and dynamics of a cold-active esterase reveals water entropy and active site accessibility as the likely drivers for cold-adaptation

**DOI:** 10.1101/2021.02.23.432564

**Authors:** Nehad Noby, Husam Sabah Auhim, Samuel Winter, Harley L. Worthy, Amira M. Embaby, Hesham Saeed, Ahmed Hussein, Christopher R Pudney, Pierre J. Rizkallah, Stephen A. Wells, D. Dafydd Jones

## Abstract

Cold-active esterases hold great potential for undertaking useful biotransformations at low temperatures. Here, we determined the structure of a cold active family IV esterase (EstN7) cloned from *Bacillus cohnii* strain N1. EstN7 is a dimer with a classical α/β hydrolase fold. It has an acidic surface that is thought to play a role in cold-adaption by retaining solvation under changed water solvent entropy at lower temperatures. The conformation of the functionally important cap region is significantly different to EstN7’s closest relatives, forming a bridge-like structure with reduced helical content providing greater access to the active site through more than one access tunnel. However, dynamics do not appear to play a major role in cold adaption. Molecular dynamics at different temperatures, rigidity analysis, normal mode analysis and geometric simulations of motion confirm the flexibility of the cap region but suggest that the rest of the protein is largely rigid. Comparison of B-factors with the closest related mesophilic and thermophilic esterases suggests the EstN7 cap region is proportionally less flexible. Rigidity analysis indicates the distribution of hydrophobic tethers is appropriate to colder conditions, where the hydrophobic effect is weaker than in mesophilic conditions due to reduced water entropy. Thus, it is likely that increased substrate accessibility and tolerance to changes in water entropy are the main drivers of EstN7’s cold adaptation rather than changes in dynamics.

## Introduction

Cold active/adapted enzymes are useful biotechnological tools due to their optimal activity at lower temperatures (1-4). Moreover, their thermolabile nature allows enzyme inactivation under mild temperature without altering product quality (5,6), such as texture and aroma in food industries. Esterase enzymes are particularly useful as they catalyse the hydrolysis of a wide variety of esters into their constituent alcohol and acid (7-11). Cold-adapted esterases are important for temperature sensitive applications such as synthesising fragile chiral compounds (12,13), cold washing laundry (14), environmental bioremediation (15) and the food industry (16).

There are several proposed molecular mechanisms by which enzymes adapt to working in cold-conditions (11,17-20). One of the most common is temperature-dependent dynamics, both globally across the whole enzyme and locally at regions important for function (19). While dynamical flux is critical to the function of general enzyme activity, cold active enzymes are thought to have evolutionarily optimized dynamics, especially around the active site. This allows the continued structural flexibility at low temperatures so enabling substrate access, binding and turnover under low energy (10,21-23). However, increased flexibility at low temperature comes with an entropic cost leading to lower overall temperature stability. In contrast, the increased structural stability at higher temperatures of mesophilic and thermophilic enzymes can come at the expense of reduced activity at lower temperature through, for example, increased rigidity (21). The features that contribute to altered dynamic properties include low levels of rigidifying residues such as prolines, glycine clustering, low salt bridge and H-bond content, less densely packed hydrophobic core and few aromatic-aromatic interactions (10,19,24,25). An alternative mechanism has been proposed in which the protein is adapted to the change in water entropy (26). As temperature drops, water molecules will become more organised and viscous so reduces the impact of the hydrophobic effect that is critical for maintaining a folded protein. Increased surface negative charge allows the protein to retain stable interactions and solvation with water under changed entropic and viscosity conditions (17,19). In some cases, the oligomerization state of an enzyme affects its flexibility and activity under low temperature conditions (27,28).

Here we determine the structure of a novel hormone sensitive lipase (HSL) family IV cold active esterase (EstN7) isolated from *Bacillus cohnii* strain N1 (29) to understand the basis of cold adaptation. *B. cohnii* was originally isolated from leather industry effluents and is considered psychrotolerant. Unlike many esterases isolated from psychrophilic organism that still have optimal activity at >20°C, EstN7 is truly cold active with optimal temperature of 5-10°C with a dramatic drop-off in activity above 20°C (29). We have previously shown that EstN7 retained function in the presence of up to 30% of various organic solvents (29), which makes it potentially useful in a variety of applications, such as fine chemical synthesis and pharmaceutical industries. The natural function of the enzyme is currently unknown and little is known about its structure; the closest structural homologue is HerE, a putative heroin esterase from *Rhodococcus* sp. strain H1 (30). The EstN7 structure revealed a protein with a highly acidic surface and a N-terminal cap region that formed a bridge-like structure allowing multiple channels for the substrate to access the active site. Analysis of the EstN7’s dynamics confirmed that the N-terminal cap was the most flexible region of enzyme in line with related esterases but the dynamic profile differed between the subunits. Overall, EstN7 was not appreciably more dynamic compared to its mesophilic and thermophilic relatives, which is counter to what might be expected for a cold-adapted enzyme.

## Results

### Structure of EstN7

EstN7 is a true cold adapted esterase with Noby et al reporting previously an optimal activity of between 5-10°C, with activity dropping of significantly at temperatures higher than 20°C (29). We determined crystal structure of EstN7 to 1.6 Å resolution (see Supporting Table S1 for statistics; PDB accession code 7b4q). EstN7 was observed as a dimer in the unit cell, which was confirmed in solution by size exclusion chromatography (Supporting Figure S1). EstN7 is a member of the hormone sensitive lipase (or IV) family with a α/β hydrolase fold (31) and consists of a central 8 stranded β-sheet surrounded by 9 helices (Figure 1a). The dimer interface area is 1049 Å^2^, equivalent to ∼8% of the average monomer accessible solvent surface area (SASA). A total of 24 residues from each monomer contribute to the dimer interface, with 13 H-bonds, 1 salt bridge and a further 16 interactions classed as salt bridges/H-bonds according to PISA (32). The two β-sheets are connected across the dimer interface via an antiparallel arrangement of strand 8 from each subunit to form effectively a single continuous sheet, with residues in the C-terminal helix and helix 8 also contributing. Each subunit is near identical with a 0.124 Å C_α_ root mean squared deviation (RMSD) and the catalytic residues occupying identical conformations (Figure S2).

**Figure 1.**
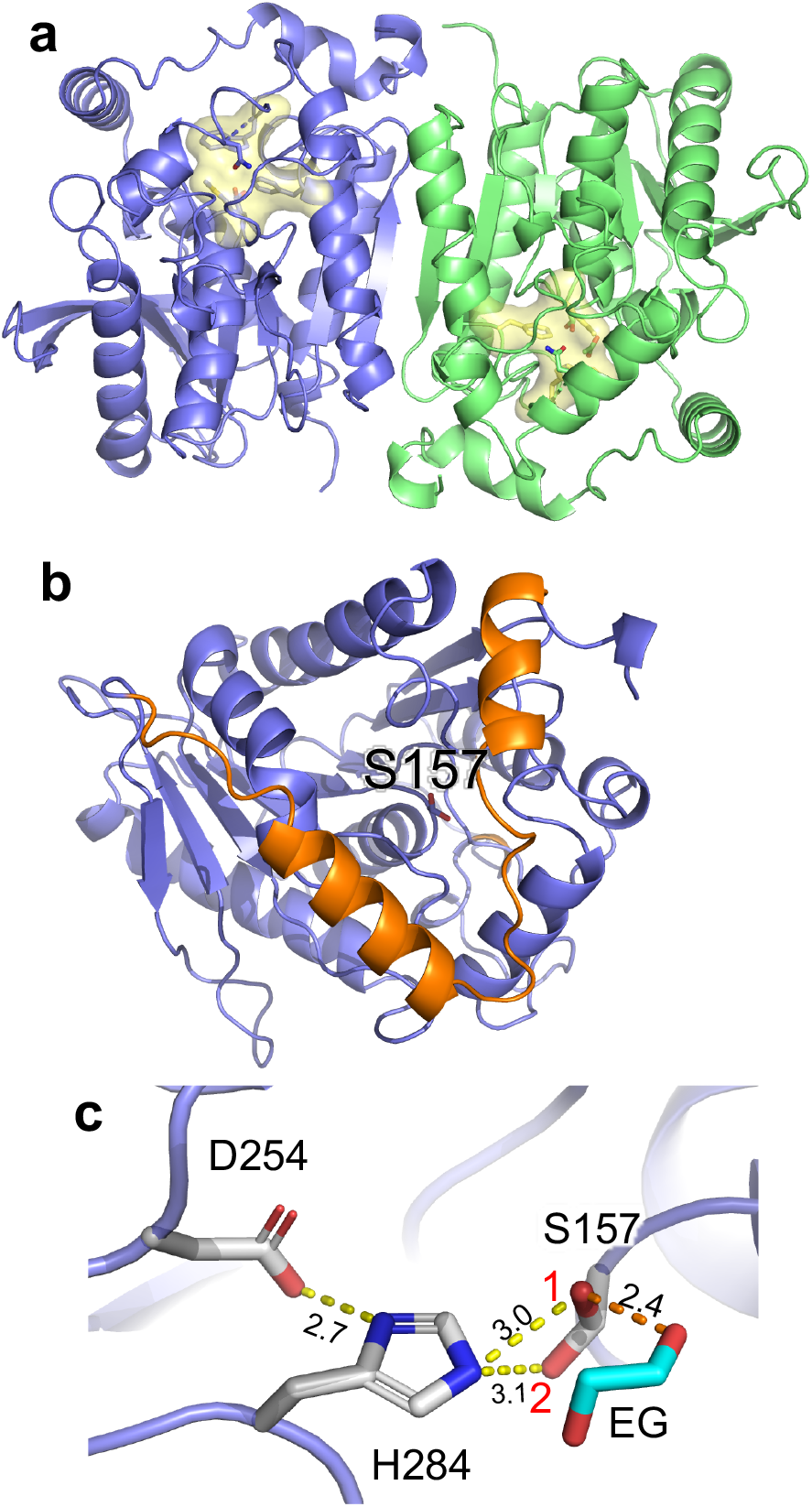
Structure of EstN7. (a) Overall structure of EstN7 with the A subunit in blue and B subunit in green. The active site is highlighted in yellow. (b) The A subunit with the N-terminal cap domain coloured orange and the nucleophilic S157 shown as sticks. (c) Catalytic triad with alternative rotamers of S157 labelled 1 and 2. An ethylene glycol (EG) is also shown as predicted to be present from electron density. Comparison with subunit B active site is shown in Figure S2a.

The nearest structural homologues are dimeric mesophilic and thermophilic bacterial esterases: a putative heroin esterase (HerE) from *Rhodococcus* sp. strain H1 (30), a mycobacterial LipW (33) and the highly enantioselective PestE from the thermophile *Pyrobaculum calidifontis* (34); the sequence identities are 40%, 37% and 35%, respectively (see Figure S3a for sequence alignment). Structural alignments generated root-mean square deviations (RMSD) of 0.62 Å, 1.08 Å and 1.18 Å, respectively, confirming conservation of structure (see Figure S3b for structural alignment). HerE and LipW represent mesophilic esterases while PestE is classed as a thermophilic esterase. An esterase from the marine bacterium *Thalassospira* sp (35) is the closest putative psychrophilic structural homologue with a sequence identity of 28% and RMSD of 1.3 Å.

Within each individual EstN7 subunit, there are 29 salt bridges according to ESBRI server (36). This is relatively high for a cold active enzyme, especially in relation to its mesophilic relatives (HerE, 21 salt bridges and LipW, 28 salt bridges). The distribution of noncovalent constraints identified by FLEXOME for rigidity analysis (37,38) is shown in Supplementary Figure S4 and is consistent with the PISA and ESBRI analyses. Multiple noncovalent constraints are visible in the dimer interface that links the two subunits. Each subunit is rich in hydrophobic tether interactions, a total of 330 such interactions being found in the dimer. Polar interactions are also common in the dimer, including 54 with effective energies of -8 to -10 kcal/mol; these are strong salt bridges.

As with other family IV esterases, EstN7 has an N-terminal cap structure. In EstN7 it comprises P7-D42 and covers the main α/β catalytic domain (Figure 1b). The cap is thought to be important for substrate access and specificity as it forms the wall of channel leading to the active site (8,10). Compared to related esterases the 2nd helical segment is shorter and further away from the 1^st^ helical segment (Figure S5). For example, in EstN7 the second helical segment is comprised of 14 residues (V24-L37) whereas for HerE it is ∼1 turn longer and is comprised of 18 residues (D25-A42), with PestE longer again comprising 21 residues (D25-N45). Analysis of EstN7 solvent accessible surface area (SASA) shows that the second helix-loop is not tightly associated with the catalytic domain as is the case in related structures but forms a bridge-like structure that reveals additional potential tunnels to the active site not present in closely related mesophilic and thermophilic esterase (Figure 2a; vide supra). The calculated volume of the active site using CASTp (39) is 1026 and 1148 Å^3^ for the A and B subunit respectively. This is significantly larger than HerE (654 Å^3^), LipW (305 Å^3^) and PestE (637 Å^3^).

**Figure 2.**
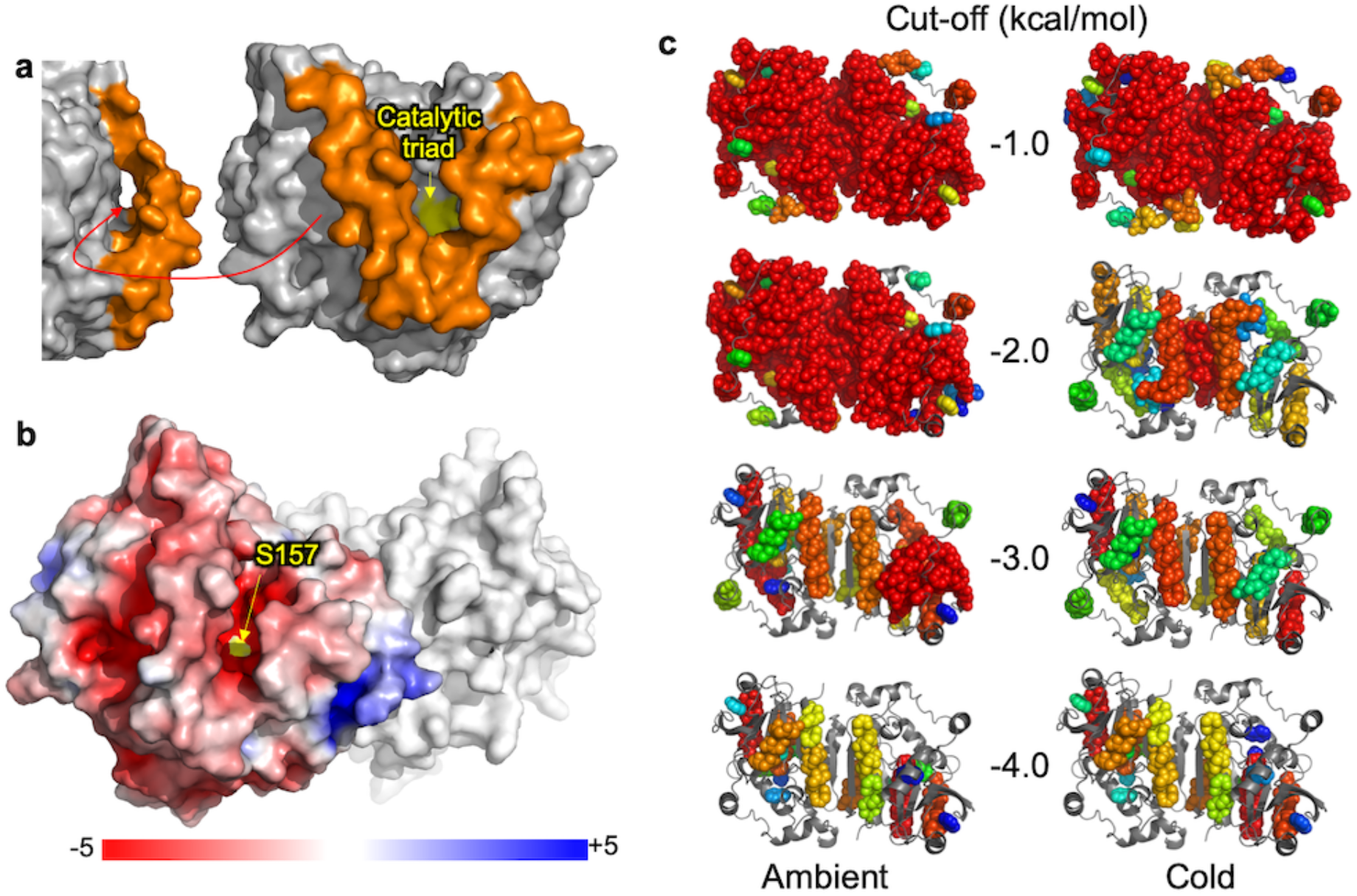
Surface and rigidity of EstN7. (a) surface view of EstN7 with the cap region coloured orange and the catalytic triad at the bottom of a deep cleft coloured yellow. Shown left is a ∼90° rotation to display a second channel. (b) electrostatic potential surface of EstN7 subunit A calculated using APBS (40). Colour scaling of electrostatic potential is shown. Rigidity features of EstN7 are shown in (c) with the 20 largest rigid clusters shown as spheres and coloured as rainbow from red (largest) to blue (20^th^ largest). The ambient column shows results using a standard analysis at cut-offs from top to bottom of -1,-2,-3 and -4 kcal/mol, while the cold column shows results when the weakening of hydrophobic tethers at low temperatures is taken into account. Flexible regions are shown as backbone cartoon and coloured grey.

The active site lies at the bottom of the cleft formed by the cap region (Figure 2a) and is comprised of Ser157, Asp254 and His284 (Figure 1c). The nucleophilic Ser157 residue is in the conserved pentapeptide motif GXSXG between strands β5 and β6 common to family IV members. The EstN7 nucleophilic serine residue resides is in the GQ**S**AG sequence motif, which would puts it in the GDSAG sequence category rather than the less populated GTSAG class (41). In both subunits, S157 adopts two distinct hydroxymethyl rotamers with ∼115° between the two (Figure 1c and Figure S2). Dual conformations of S157 have been observed previously for related esterases with the different rotamers thought to be representative of the catalytic cycle rather than an artifact of crystallisation (30,33,42); one rotamer represents the hydrolytic competent state while the other facilitates product release and prevents reverse reactions. Both rotamers are capable of polar interactions with His284 with distances of 3.0 Å and 3.1 Å. Rotamer 1 faces towards the acyl binding pocket as expected for its role in catalysis so is likely to represent the hydrolytically competent state. Electron density that equates to ethylene glycol was also observed in the active site of both subunits, with an additional ethylene glycol found in subunit B (Figure S2b). Rotamer 1 is closest to the observed ethylene glycol moiety bound in the active site within the alcohol binding pocket, with the hydroxyl group within H-bonding distance (2.4Å) of the alcohol group (Figure 1c). Ethylene glycol could thus act as mimic for the alcohol product/substrate associated with esterase activity. The remaining residues comprising the catalytic triad, Asp254 and His 284, are found between β7-α8, and β8-α9, respectively. The oxyanion hole, which helps stabilise intermediates during catalysis, is located close by and is comprised of the conserved 83-HGGG-86 motif (Figure S2a).

Sequence analysis of EstN7 reveals a high proportion of charged residues, with acidic residues dominating (Asp-Glu:Lys-Arg ratio being 48 to 33). Surface electrostatics were calculated using APBS (Adaptive Poisson-Boltzmann Solver) (40) and revealed the surface of EstN7 is largely negative with a few basic patches (Figure 2b). The substrate binding cavity is largely acidic in nature (Figure 2b). The thermophilic PestE has a lower number of charged residues (68 *versus* 88 for EstN7) with a nearly 1:1 ratio of basic and acidic residues (Asp-Glu:Lys-Arg ratio being 35 to 33), which is manifested in its electrostatic surface profile (Supporting Figure S6).

### Structural rigidity of EstN7

Rigidity dilution (43) provides information on the relative rigidity/flexibility of different parts of a protein crystal structure. In this process, polar constraints are gradually eliminated in order of strength by lowering a “cut-off” value which excludes weaker bonds from the analysis. Flexible loop regions lose rigidity at small cut-offs whereas the “folding core” of the protein retains rigidity longest. Previous studies on the rigidity of extremophiles (38,44,45) have been consistent with the “corresponding states” hypothesis(46); thermophilic and hyperthermophilic enzymes are more rigid than their mesophilic counterparts at a given cut-off, and the rigidity of a thermophile at a large cut-off resembles that of a mesophile at a smaller cut-off. We therefore assessed the rigidity of EstN7 using pebble-game rigidity analyses at a series of cut-off values (−1, -2, -3 and -4.0 kcal/mol). The resulting rigid cluster decompositions are shown in Figure 2c.

On carrying out a conventional rigidity analysis under ambient conditions (the default for mesophilic proteins), the EstN7 structure in fact appears very rigid (Figure 2c, “ambient” column). At cut-offs of -1.0 and -2.0 kcal/mol, a single large rigid cluster extends across most of the body of the dimer, with only the cap regions appearing independently flexible. Even at a cut-off of -3.0 kcal/mol a large rigid cluster spanning a significant portion of chain B is visible in the rigid cluster decomposition. By -4.0 kcal/mol cut-off, only individual helices appear as large rigid clusters. This apparently enhanced rigidity in a cold-adapted enzyme is counter-intuitive. However, a previous study including a citrate synthase from an Antarctic microorganism (44) shows how to resolve the discrepancy. In the conventional analysis a hydrophobic tether removes two degrees of freedom from the molecule. To reflect the weakening of the hydrophobic effect at lower temperature, a second analysis in which each hydrophobic tether removes only one degree of freedom (Figure 2c, “cold” column) gives more appropriate results for a cold-adapted enzyme. While a single large rigid cluster extends across most of the dimer at the smallest cut-off of -1.0 kcal/mol, at -2.0 kcal/mol much of the structure is flexible, with individual rigid helices, and the remaining largest rigid cluster is found at the centre of the dimer, spanning the dimer interface. Both “ambient” and “cold” analyses concur that the cap regions are the most flexible parts of the EntN7 structure.

Comparative rigidity analysis with HerE, LipW and PestE is shown in Supporting Figure S7. For the thermophilic PestE, the monolithic rigidity observed at -1 kcal/mol cut-off is also observed at -2 kcal/mol, which is indicative of greater structural rigidity normally considered important for stability at high temperatures. For mesophilic HerE and LipW, the rigidity profile at the -1 kcal/mol cut-off, with even weak polar interactions included as constraints, gives a monolithic rigid cluster that extends across the entire structure. At the -2 kcal/mol cut-off, there is still a large rigid cluster across the structure, but more regions of flexibility are now observed, including around the active site cleft (Figure S7). Thus, rigidity analysis indicates that the mesophilic homologues are less rigid than the thermophilic PestE, as expected. The rigidity of EstN7 appears comparable to that of PestE in an ‘ambient’ analysis, as both structures are rich in salt bridge interactions. However, a ‘cold’ analysis of EstN7, allowing for the weakening of the hydrophobic effect at low temperatures (Fig. 2c), indicates that the structure is less rigid than the mesophilic or thermophilic homologues.

### EstN7 dynamics

Using a combination of B-factors and temperature-dependent molecular dynamics (MD), flexible regions of EstN7 were identified. In line with the rigidity analysis, both MD simulations and B-factors show clearly that the cap region is the most dynamic aspect of EstN7 (Figure 3a-c), as is the case with esterases in general (7,10,31). However, the cap region does appear to have different dynamics between the two subunits, including at different temperatures (Figure 3b-c and Figure S8). While both subunits have the bridge-like structure associated with the second helical segment, the cap region is marginally closer to the catalytic domain in subunit A compared to B (Figure 3a). In subunit A the loop immediately following the second helix has the highest B-factors whereas it is C-terminal half of the second helix for subunit B (Figure 3d). In subunit A, a second region comprising residues 70-75 adjacent to the C-terminal end of the cap region display higher than average B-factors.

**Figure 3.**
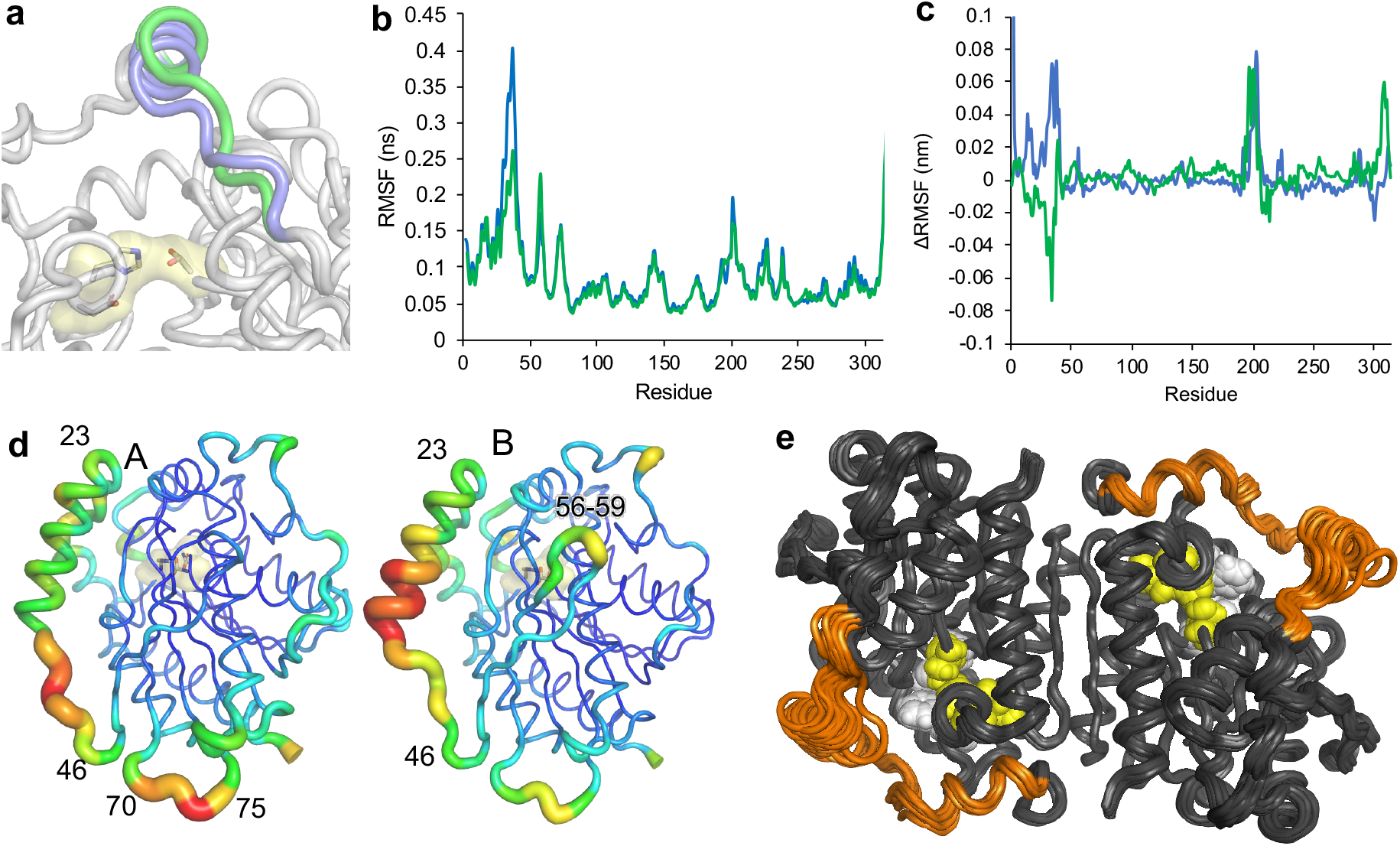
EstN7 dynamics. (a) Comparison of the conformation of the second helical component of the cap region in the A (blue) and B (green) subunit. The catalytic triad is shown as sticks for reference. (b) Residue-by-residue root mean squared fluctuation (RMSF) of Cα during 100 ns molecular dynamic simulations. Overlayed are the values for A (blue) and B (green) subunit. RMSF is reported as an average of 5 independent simulations run for 100 ns at 298K (25°C). (c) RMSF difference between simulations run at 283K and 308K for the A and B subunit. The ΔRMSF was calculated by subtracting 283K value from the 308K value. A positive or negative value reflects increased RMSF either higher or lower temperature, respectively. The RMSF plots at each temperature are shown in Figure S8. (d) Putty diagrams of the A (left) and B (right) subunits to represent the B-factor changes over the backbone. Higher values correspond with thicker, redder putty. B-factor values over the cap region are shown in Figure 4. (e) An ensemble of twenty structures generated by geometric simulations of flexible motion biased parallel and antiparallel to the ten lowest-frequency nontrivial normal modes, representing the amplitude of flexible motion in the structure achievable while maintaining its bonding and steric geometry. The catalytic triad is shown as yellow spheres and the plug residues as white, while the cap region is coloured orange. The greatest extent of conformational variation is seen in the cap regions, which can freely explore motions of several Å in amplitude.

A similar trend was observed for MD simulations of EstN7 run at 298K (Figure 3b). The root mean squared fluctuations (RMSF) over the C_α_ atoms revealed that residues within the cap region are flexible in both subunits together with the adjacent 70-75 loop. The MD simulations suggests that the A subunit is more dynamic in the cap region around the second helical segment, which could imply this helical region has more scope for movement to potentially a more open conformation. MD also shows that the turn comprising residues 56-59 also has a higher dynamic profile, which is backed up by the increased B-factors in the equivalent region in subunit B (Figure 3b-c). MD also suggests that a short region centred on residue 201, which is adjacent to the cap domain and close to the N-terminal (Figure S8c), is also flexible but B-factors indicate this region is not overly dynamic. Simulations run at 283K and 308K (10°C and 35°C, respectively) show that the majority of the protein does not undergo any significant change in flexibility (Figure 3c and Figure S8a-b). The short segment centred on residue 201 does appear to be more flexible at higher temperatures. As with the B-factors and 298K MD simulation, the temperature-dependence of the cap region differs depending on the subunit (Figure 3c). The cap region of the A subunit becomes more dynamic as at the higher temperature whereas the B subunit is slightly less dynamic. Comparison of the RMSF data at each temperature (Figure S8a-b) suggests that at 283K, the cap regions have similar dynamics regimes compared to the different regimes observed at higher temperatures.

Cold active enzymes are generally considered more dynamic, especially around regions important for function compared to their mesophilic and thermophilic counterparts (19). We compared the general trend in C_α_ B-factors observed across the cap region for the A subunit of EstN7 with those found for the A subunits of HerE (30) and the thermophilic PestE (34) (Figure 4). Overall B-factors for the cap regions for each of the esterases are higher than the average backbone value. EstN7 has an overall average B-factor ∼26 Å^2^, which is higher than both PestE and HerE (both ∼18 Å^2^). However, the change in the normalised B-factor for the cap region is lowest for EstN7 (1.6-fold) compared to PestE (1.9-fold) and HerE (2.0-fold). Thus, it appears that the cap domain of EstN7 is proportionately less flexible compared to even thermophilic relatives. In comparison with EstN7, the highest B-factor values for both HerE and PestE are in the first helical region of the cap structure (Figure 4), with residues 15 to 19 missing from the structure of PestE presumably due to high mobility.

**Figure 4.**
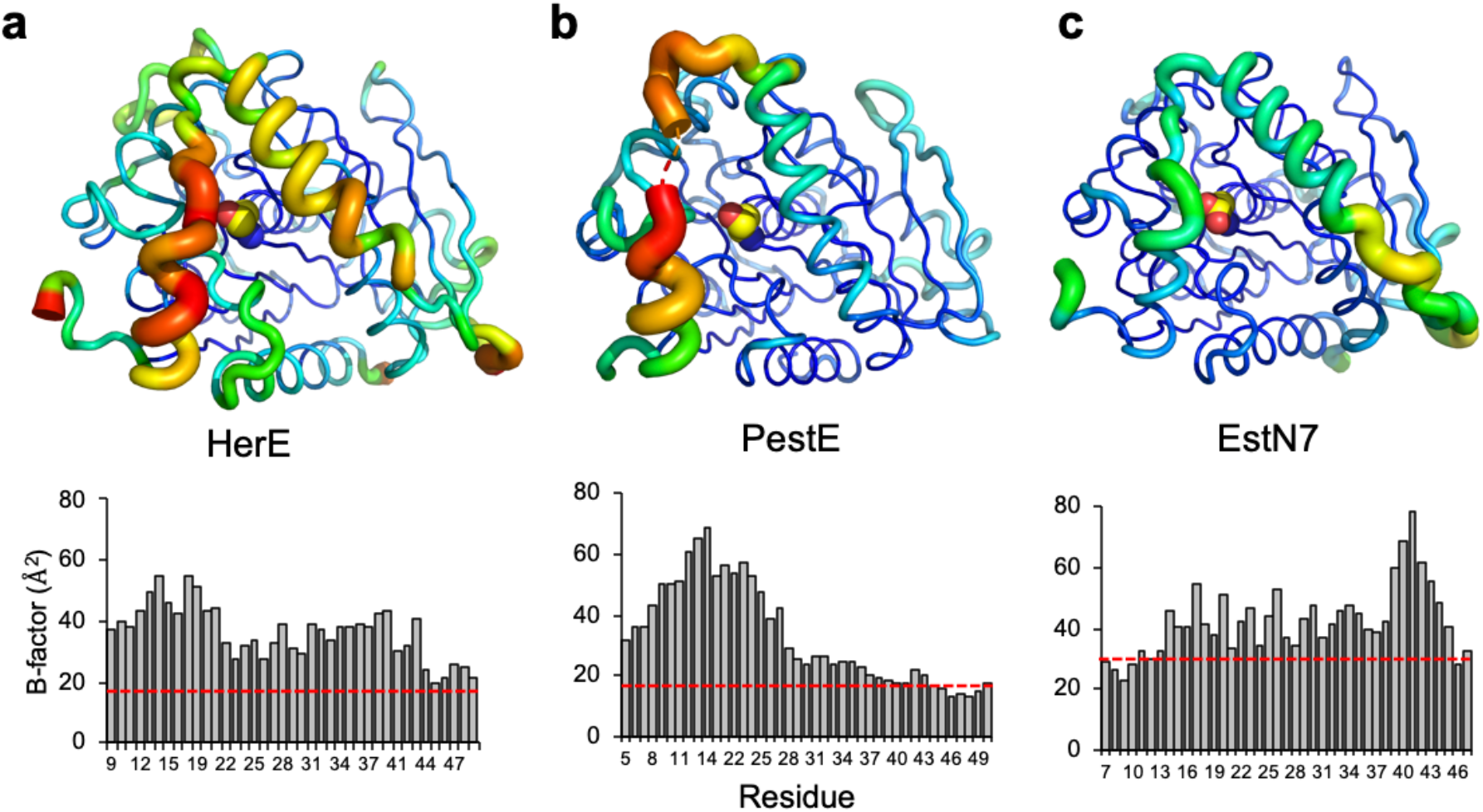
B-factor analysis of HerE and PestE. Relative changes in B-factor along the backbone for (a) HerE (PDB 1lzk) (30), (b) PestE (PDB 2yh2) (34) and (c) EstN7 (PDB 7b4q). Top panels are the putty diagram of the A subunit of each esterase. Putty diagrams were generated in PyMOL. The same scaling is applied across the 3 structures so represents a comparative analysis. The lower panel represents the Cα B-factor values for the cap region. The average Cα B-factor across the whole structure is shown as a red line (HerE, 18 Å^2^; PestE, 18 Å^2^; EstN7, 26 Å^2^). Residues 15 to 19 are missing from the PestE plot as they are absent in the PDB file.

B-factors provide an experimental measure of local thermal fluctuations in a crystal structure, while MD simulations over nanosecond timescales probe the capacity for concerted local flexible motion. Geometric simulations of flexible motion (44,47,48) provide a complementary probe of the global, collective flexible motion achievable on longer timescales. In this approach, we first obtain low-frequency normal modes of motion for the protein, representing “easy” directions of motion which the protein can explore with minimal restoring force or energetic penalty. We then explore motion of the structure along these directions while using constraints to retain the bonding geometry of the starting structure. We therefore carried out elastic network modelling of EstN7 on a one-node-per-residue basis with a spring interaction range of 10.0Å, extracting the six trivial rigid body motions and the ten lowest-frequency nontrivial normal modes (modes 07 to 16). Examination of the resulting eigenvectors showed that the two lowest-frequency nontrivial motions, modes 07 and 08, are collective twisting and bending motions of the entire dimer, while modes 09,10 and higher are more localised on flexible loop regions and in particular on the cap regions. Geometric simulations biased parallel and antiparallel to each mode eigenvectors then generated an ensemble of flexible variations on the EstN7 structure, representing the amplitudes of motion achievable in these “easy” directions of low-frequency motion while maintaining steric exclusion and the covalent and noncovalent bonding geometry of the crystal structure. This ensemble is shown in Figure 3d. The capacity of the entire dimer for collective motion is very limited by the well-constrained dimer interface, with the central part of the dimer displaying little variation in the ensemble. In contrast, the cap regions clearly display a capacity for large-amplitude flexible motions which move the cap through amplitudes of several Ångstroms over the active site cleft below. Two low-frequency modes, 09 and 10, are particularly localised onto the cap regions; a visualisation of these motions, and the consequent variations in the primary and secondary channel geometries, are shown in Supplementary Figure S9. The flexible dynamics of the cap regions seen in the geometric simulations are consistent with the hypothesis that conformational flexibility in this region is a key element of the cold adaptation of EstN7, offering greater access to the active site.

### Thermal stability of EstN7

Circular dichroism was used to assess the structural thermal stability of EstN7. At 15°C, EstN7 had a significant helical character with characteristic troughs at 210 and 222 nm suggesting the enzyme is largely folded; these troughs become shallower above 15°C but still suggest a largely fold enzyme up to 50°C (Figure 5a). Between 50-60°C, EstN7 undergoes a major structural transition, with the enzyme largely unfolded at 60°C giving the CD spectra shown in Figure 5a. Thermal unfolding of EstN7 monitored by CD at 208 nm revealed a major transition between 35-55°C, with a T_m_ of 51°C (Figure 5b).

**Figure 5.**
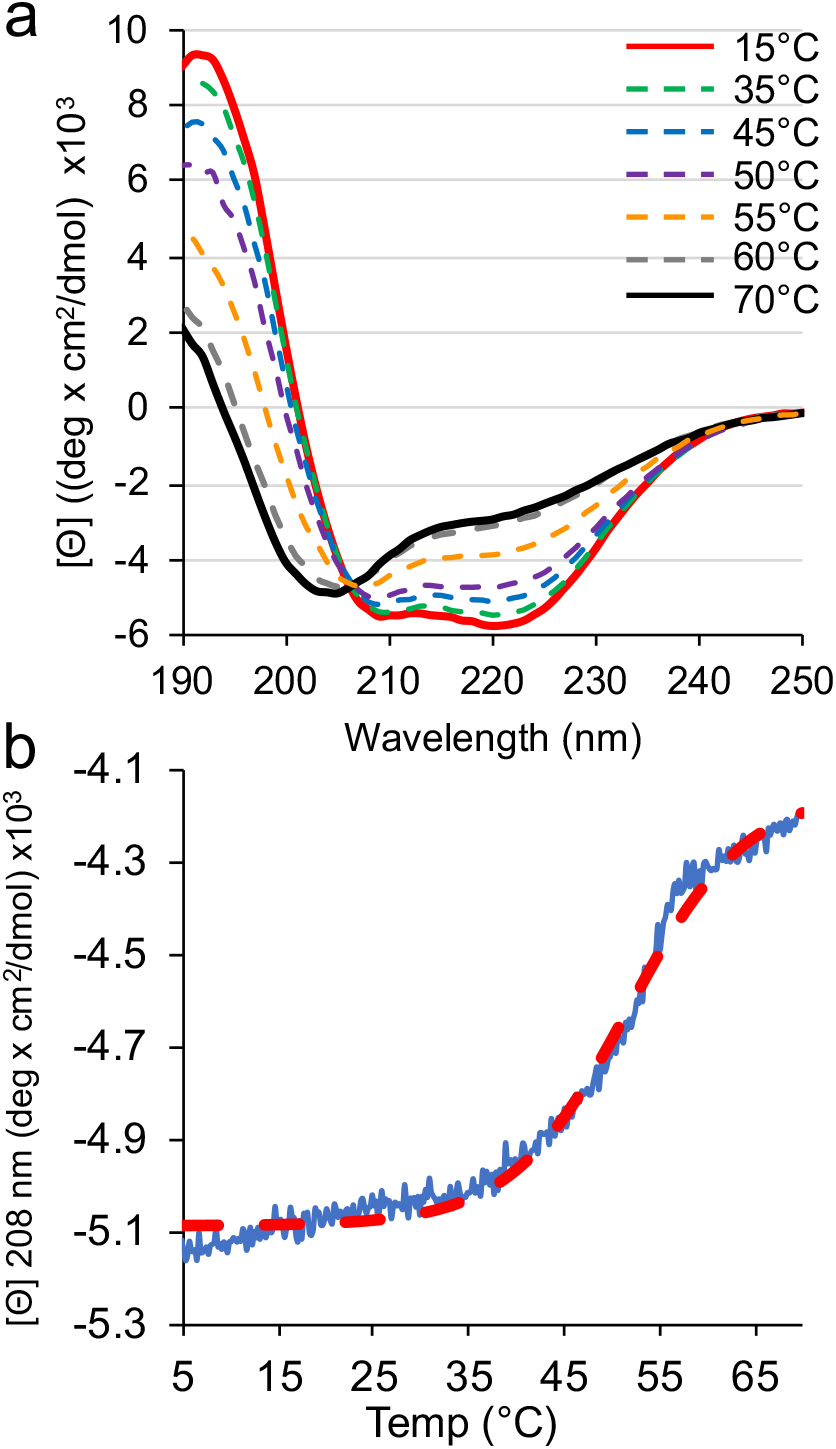
Thermal stability of EstN7 monitored by CD spectroscopy. (a) CD spectra of EstN7 at 15°C (red solid line), 35°C (green dashed), 45°C (blue dashed), 50°C (purple dashed), 55°C (orange dashed), 60°C (grey dashed) and 70°C (black solid line). (b) Temperature-dependent changes in absorbance at 208 nm for EstN7. The red line is a Boltzmann sigmoidal curve fit to the major transition during temperature ramping.

## Discussion

Cold-active esterases represent an important technical enzyme class as well as providing fundamental insights into adaptation of the enzyme structure-function relationship for low-temperature activity. Cold-active esterases are particularly important for industrial processes that require low temperatures such as food processing, low temperature laundry, bioremediation and fine chemical production (7,9-11,25). Thus, it is important to understand the underlying structural mechanism by which cold adaption takes place. Here we have determined the crystal structure of EstN7 from *B. cohnii*, a cold active esterase belonging to the hormone sensitive lipase IV family (Figure 1). The nearest structural homologues are <40% identical so the structure provides new insights into this important enzyme class. EstN7 is characteristic of the family IV esterase, including the classical α/β hydrolyase dimeric fold with the extended β-strand linking the two monomers and the presence of a N-terminal cap region (Figure 1). The catalytic triad lies at the bottom of a cleft as with related family IV esterases(8,25), with the cap region contributing to the walls of the cleft (Figure 2a). However, the nature of the cap region is significantly different compared to its closest relatives, as discussed below.

The question thus arises as to the molecular basis for cold adaptation. EstN7 can be considered a true cold active esterase based on its temperature activity optimum (29). Many esterases are termed psychrophilic due to the organisms they derive from rather than their inherent cold-optimised activity, with the enzymes themselves commonly being mesophilic with regards to their structural stability and activity (10,35,49). Previous work showed that EstN7 has optimum activity at 5-10°C, dropping off significantly between 20-30°C (29). Thermal unfolding however suggests that EstN7 is relatively stable (Figure 5; T_m_ =51°C). It is generally thought that cold-active enzymes tend to be thermally unstable, losing structure as well as function at relatively low temperatures primarily through increased dynamics. However growing evidence suggests that this does not always hold true as shown here and elsewhere (17,23,28,50-52). Thus, at least for these psychrophilic enzymes, global stability is not a physico-chemical restriction on cold activity. This in turn raises the question of the role of dynamics as a main driver for cold adaptation, which is discussed below.

Various structural mechanisms are suggested to play a role in cold-adaptation (18,19,24). Water entropy is key to protein thermodynamic stability, especially the hydrophobic effect, and its loss is important to the phenomenon of cold-denaturation. This is due to the change in water organisation and viscosity at low temperatures and so there is a greater energetic cost to breaking the solvent H-bond network (26). Protein surface charge is thought to play a role in addressing water entropy (19), with EstN7 having a high proportion of charged residues (25% of total residues are Arg, Asp, Glu or Lys), and an acidic surface in keeping with its high Asp/Glu:Lys/Arg ratio (Figure 2b). Rigidity analysis of the EstN7 structure under “cold” conditions (through weakening hydrophobic interactions to mimic the reduced hydrophobic effect due to lower water entropy) provides further evidence of a water entropy driven mechanism (Figure 2b). A negatively charged surface can offset water entropy by maintaining the critical solvation layer through favourable interactions with the water. Indeed, EstN7 is known to be tolerant to high concentrations of organic solvent (29); the retention of a water shell may not only be advantageous for cold-adaptation but may also be useful for solvent tolerance. Local high concentrations of charge could lead to charge repulsion and thus increased flexibility but both B-factors and MD simulations do not suggest this is the case here (Figure 3); any charge repulsion can potentially be offset through interactions with water. It has been suggested that charged surfaces play a role in halo-tolerance (53) but EstN7 does not display any obvious salt tolerance (Figure S10).

As mentioned above, protein dynamics are also thought to play a critical role in cold-adaptation. While a commonly perceived idea is that cold-adapted enzymes are inherently more flexible, it may not always hold true (54); this appears to be the case here given the inherent dynamics of EstN7 *per se* and when compared to its mesophilic and thermophilic relatives. EstN7 does not have a particularly low level of rigidifying residues, such as prolines (55), with, for example, EstN7 having more proline residues (24) than the thermophilic PestE (20). Increased glycine content and clusters are also thought to increase flexibility; EstN7 and PestE have a comparable number of glycines (24 and 23, respectively), with the only glycine clusters associated with the conserved glycine rich motifs of the oxyanion hole and nucleophilic serine. EstN7 also has a substantial number of salt bridges, with the mesophilic HerE having a lower number (29 *versus* 21, respectively). The hydrophobic cores of cold-adapted enzymes are also thought to be less well packed (19,24); rigidity analysis shows that EstN7 has a well packed core rich in hydrophobic tether interactions. Indeed, EstN7 has a similar rigidity to the thermophilic PestE under ambient conditions, and more rigid than mesophilic relatives (Figure S7). MD analysis at 283K and 308K suggests that most of the residues do not significantly change in terms of their dynamic profile (Figure 3c).

Changes in local dynamics, especially in regions important for catalysis, are also thought to be important for cold adaptation (17,21,23,56). For example, while B-factors have been reported to be generally lower for cold-adapted proteins, these increase locally around regions important for catalysis (57). The cap region is a feature common to many esterases and dynamics of this structure are important for activity (10). In EstN7, the 2nd helical component of the cap region is shorter than its closest thermophilic relative and has less helical content (Figure 1b and Figure S5). In line with related esterases, B-factors (Figure 3d), rigidity analysis (Figure 2c), geometric simulations (Figure 3e) and molecular dynamics (Figure 3b-c) show that EstN7 cap region is the most conformationally flexible part of EstN7 but the rest of the protein remains relatively rigid. Comparison of B-factors does not suggest it is especially more dynamic compared to closely related esterases (Figure 4). Moreover, relative with the average B-factor over the whole protein, the cap region of EstN7 is comparatively less dynamic than two of its relatives. Thus, it does not appear that dynamics of the functionally important cap region is significantly different to its closely related mesophilic and thermophilic structural relatives. MD simulations reveal that the dynamics of the cap region differs between the EstN7 subunits (Figures 3b-d and Figure S8). At higher temperatures subunit A has an inherently more flexible cap region than B (Figure 3b-d). Furthermore, relative change in dynamic profile of the cap region as a function of temperature is different in each subunit (Figure 3c) and suggest that they coalesce to a similar profile in both subunits at 10°C (Figure S8).

The results of rigidity analysis and of geometric simulations of flexible motion are consistent with the above hypotheses. The rigidity analysis suggests that EstN7 at lower temperatures has rigidity comparable to mesophilic enzymes at ambient temperatures when the weakening of the hydrophobic interactions at low temperature is considered (Figure 2c and Figure S7). During flexible motion along low-frequency normal mode directions the cap regions are capable of substantial amplitudes of motion while retaining the local geometry of the crystal structure (Figure 3e); for example, the alpha-helical portion of each cap remains well constrained by backbone hydrogen bonds and moves as a unit.

An alternative mechanism for cold activity could be the conformation of the cap regions rather than just its dynamics. Analysis of the protein’s surface together with channel prediction suggests that the cap is partially detached from the main catalytic domain forming a bridge-like structure, with channels to the active site present from both sides (Figures 3a and 6). These secondary tunnels are absent in related esterases (Figure 6) despite the cap region following a similar path. The conformation of the cap domain in turn results in a wider main channel and the generation of additional tunnels to access the active site. In contrast, there is only one clear, common channel into the active site for the closely related structural homologues (Figure 6b-c). As well as increased access routes, the active site cavity volume is > 1.7 fold larger for EstN7 than its closest mesophilic and thermophilic relatives. It has been proposed that increased cavity size is a mechanism for cold adaption and can promote substrate binding even at low temperature (28,59). Overall, our findings are consistent with the hypothesis that the bridge-like structure of the cap region grants greater access to the active site than in related esterases.

**Figure 6.**
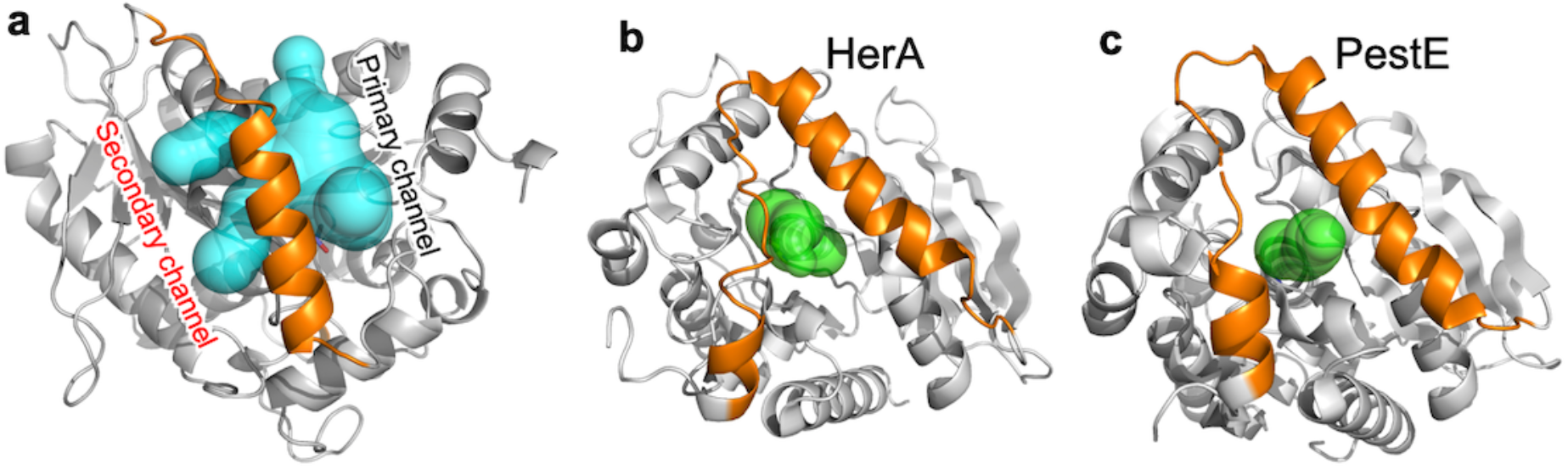
Available tunnels for access to the active site of (a) EstN7, (b) HerA and (c) PestE. Tunnels were calculated using CAVER 3.0(58) with minimum probe radius set to 1.4 Å.

## Conclusion

Cold active esterases provide an insight into the molecular mechanism by which enzymes evolve to work in extreme conditions. EstN7 represents a novel cold-active esterase, with its nearest structural homologue having 40% or less sequence identity. A highly acidic surface profile helps retain structure when water becomes more viscous at lower temperatures. However, one perceived common feature of cold adapted enzymes does not appear to be critical here: higher local and global dynamics compared to mesophilic and thermophilic counterparts. Thus, EstN7’s prime cold-adaptation mechanism is likely to be dealing with change in water entropy and allowing improved access for substrate to the active site. In enzymes (like esterases) with the active site located at the end of molecular tunnels, the substrate must firstly navigate the channel. Flexibility around the cap regions comprising the channel is common in both mesophilic and thermophilic esterases. EstN7 has relatively similar and to some extent lower dynamics in this region. However, the cap region is not fully associated with the main catalytic domain as it forms a bridge structure so allowing multiple channels for substrate to access the active site. Whether these combined features of cold-adaptation are common amongst esterase and enzymes in general remains to be seen but it does provide potential protein engineering routes to improve activity at lower temperatures for biotechnological applications.

## Experimental procedures

### Protein production and purification

Isolation of EstN7 encoding gene from *Bacillus cohnii* strain N1 and characterization of the recombinant enzyme were described previously (29). EstN7 present within the pET28 a(+) plasmid was used to transform *E. coli* BL21 DE3 cells. Single colonies where selected from LB agar plates supplemented with kanamycin and cultures left overnight in LB medium, which was used to inoculate 1L fresh LB medium containing 35µg/ml kanamycin. The culture was incubated at 37°C until reaching an A_600_ of 0.6 upon which EstN7 production was induced by the addition of isopropyl-β-dthiogalactopyranoside (IPTG) to a final concentration of 1mM; the culture was incubated for a further 18h at 37°C. Cells were then harvested by centrifugation at 5000x *g* for 20 minutes and the pellets washed and re-suspended in 50 mM Tris-HCl buffer, pH 8.0 (Buffer A). Cells were lysed by sonication for 10 min (8 cycles of 30 s, 18,000 Hz each with interval of 1 min on ice between cycles). Cell lysate was further clarified by through centrifugation at10,000 x *g* for 10 minutes. Ni^2+^ affinity chromatography was used initially to purify recombinant EstN7, the cell lysate was separated from the cell debris. The crude mixture was loaded onto a nickel affinity chromatographic gel column (Sigma-Aldrich Co) and the column washed with buffer A containing 20 mM imidazole. EstN7 was eluted using buffer A supplemented with 500 mM imidazole. Samples where further purified using a Superdex 200 HiLoad 26/600 size exclusion column (Cytiva). The proteins were eluted using buffer A at room temperature.

### Structure determination

Crystallization conditions were explored using PACT premier™ and JSCG screens. The trials were performed in 96 well plates with a crystallization robot using sitting drop vapor diffusion method. For each trial, 1μL of the protein sample (7mg/mL) and reservoir solution were mixed and equilibrated against 30 μL of the reservoir solution. The plates were incubated at 20°C and checked regularly for crystals formation. EstN7 crystals were obtained after 5 days in several crystallization conditions. The crystal producing the best diffraction was grown in 0.2M lithium chloride, 0.1M sodium acetate, 20% (w/v) PEG 6000, pH 5.0 (JCSG screen condition A05). Crystals were harvested by looping into thin nylon loops and plunged into liquid N_2_, for transfer to Diamond Light Source, Harwell, UK. Data were collected at beamline I03. Diffraction data were reduced with the XIA2 package using DIALS software. Scaling and merging were completed with AIMLESS and CTRUNCATE, part of the CCP4 package. The structure was solved with molecular replacement using PHASER(60), with HerE (1lzk.pdb (30)) as a starting model. Refinement with REFMAC (61,62) was alternated with graphics sessions in COOT (63) till convergence. Channels were identified using CAVER 3.0 (58) with a minimum probe radius of 1.4 Å and maximum distance of 10 Å.

### Rigidity analysis, normal mode analysis, and geometric simulations of flexible motion

Pebble-game rigidity analysis(37) (in which degrees of freedom are matched against constraints in a directed graph representing the covalent and noncovalent bonding network of the protein, identifying rigid clusters and flexible regions in the structure), elastic-network normal mode analysis(64) (identifying directions of low-frequency flexible motion), and geometric simulations of flexible motion(47,48) (in which the structure is moved along a normal mode direction while maintaining its bonding geometry and steric exclusion) were carried out using a novel software implementation (FLEXOME) recently written by one of us (Wells). FLEXOME implements a body-bar version of pebble-game rigidity analysis which duplicates the functionality of the older FIRST software(37) and includes recently identified corrections to the handling of short salt bridge interactions(38). The elastic network utility of FLEXOME provides the same functionality as standard codes such as Elnemo(64), but makes use of inverse iteration to extract only a limited number of the lowest frequency modes rather than fully inverting the Hessian matrix. The geometric simulation utility of FLEXOME replicates the functionality of the FRODA(48) module, which was originally implemented within FIRST, and is written to explore flexible motion of the all-atom protein structure along low-frequency normal mode directions as has been previously done using the combination of FIRST/FRODA and Elnemo(44,47). FLEXOME is available to researchers upon request from the repository at Bath (https://doi.org/10.15125/BATH-00940) subject to non-commercial licence conditions.

In this instance, hydrophobic tethers and noncovalent polar interactions were identified by geometric criteria, and polar interactions were assigned effective energies from -0.0 to -10.0 kcal/mol based on the donor-hydrogen-acceptor geometry(38). Rigidity analysis was then carried out using a progressively lowered cutoff to eliminate weaker hydrogen bonds, giving a “rigidity dilution”(43) revealing the relative rigidity of different portions of the structure. In a standard analysis, polar interactions introduce five constraint “bars” into the protein and hydrophobic tethers introduce two bars; each bar removes a degree of freedom from the system. In this case we carried out both the standard analysis and a variant appropriate to cold conditions(44), in which hydrophobic tethers introduce only one “bar”.

The elastic network model was created by extracting one node per residue, using the Cα positions, and assigning springs of uniform strength between all pairs of nodes separated by a distance of less than 10Å. Six rigid-unit motions of the structure were constructed analytically by FLEXOME and the ten lowest-frequency nontrivial modes extracted by inverse iteration; these are denoted as modes 07 to 16.

Geometric simulations were carried out retaining the noncovalent constraint network selected by a cut-off value of -4.0 kcal/mol, exploring motions parallel and antiparallel to each of modes 07 to 16. The bias in each step of the simulation was given by the (normalised) mode eigenvector multiplied by a scale factor of 0.1Å; up to 500 steps were carried out in each simulation, with every 50^th^ step saved in PDB format as a “frame”. The tolerance criterion on deviations from the bonding and steric geometry was 0.1Å; each simulation halted after 500 steps or when the geometry could no longer be restored to within tolerances, which typically occurred within ∼100-200 steps. The rigidity analysis, elastic network modelling and geometric simulation approaches are all rapid and computationally lightweight(47); all of the above modelling was carried out on a laptop computer in a few CPU-hours.

### Molecular dynamics simulations

Molecular dynamics on the dimeric EstN7 crystal structure were performed using Gromacs 4.6 (65) on the Hawk Supercomputing resource at Cardiff University (part of Supercomputing Wales). The pdb was initially converted to gmx format using the AMBER99SB forcefield (66). The protein was then placed centrally in a cubic box at least 1 nm from the box edge. The protein system was then solvated with water molecules and total charge balanced to zero with Na+ and Cl-ions. The protein was then energy minimised to below 1000 kJ/mol/nm with an energy step size of 0.01 over a maximum of 50000 steps. The system was then temperature and pressure equilibrated before molecular dynamics runs at 283K, 298K or 308K and 1 atmosphere pressure for 100 ns. Four independent MD simulations were chosen at each temperature for further analysis (see Figure S11 for RMSD over the course of the simulation for each production MD run) with the Cα residue-by-residue root mean squared deviation fluctuation (RMSF) reported as an average of the four simulations.

### Circular dichroism spectroscopy

Immediately prior to analysis, lyophilised EstN7 freeze dried in 50 mM potassium phosphate buffer, pH 7, was resuspended in water to 5 µM. Circular dichroism spectra were measured using 5 µM protein in approximately 5 mM potassium phosphate buffer, pH 7. Measurements were obtained at Bath University School of Chemistry on Chirascan CD Spectrophotometer (Photo Physics, Surrey, UK), using 0.1 mm pathlength quartz cuvette after blanking with 5 mM Tris pH 8.0. Absorbance was measured between 190 and 260 nm, at 1 nm intervals with a scan rate of 7.5 nm/min. Full spectra were recorded at 5°C intervals starting at 15°C and ending at 70°C. The thermal melt (*T*_m_) was determined by increasing temperature at a ramp rate of 1°C per minute using a Quantum Northwest Peltier and polarized absorbance at 208 nm was recorded at every 0.2°C change up to 70°C. Data obtained was converted to molar ellipticity and fitted to a Boltzmann sigmoidal curve in Origin V 2020 (MA, USA).

## Data availability

The structure of EstN7 has been deposited in the PDB under accession code 7b4q. FLEXOME is available to researchers upon request from the repository at Bath (https://doi.org/10.15125/BATH-00940) subject to non-commercial licence conditions. All other data is available on request from the authors.

## Supporting Information

This article contains supporting information: Supporting Table 1, Supporting Figure S1-S11.

## Acknowledgements

We would like to thank the staff at the Diamond Light Source (Harwell, UK) for the supply of facilities and beam time, especially Beamline I03 staff. We would also like to thank the Protein Technology Hub facility in the School of Biosciences, Cardiff University for access to protein purification and analysis facilities. Molecular dynamic simulations where run on the Hawk facility as part of Supercomputing Wales under project code scw1631.

## References

1. Sarmiento, F., Peralta, R., and Blamey, J. M. (2015) Cold and hot extremozymes: industrial relevance and current trends. Frontiers in bioengineering and biotechnology 3, 148

2. Al-Ghanayem, A. A., and Joseph, B. (2020) Current prospective in using cold-active enzymes as eco-friendly detergent additive. Applied Microbiology and Biotechnology 104, 2871–2882

3. Al-Maqtari, Q. A., Waleed, A.-A., and Mahdi, A. A. (2019) Cold-active enzymes and their applications in industrial fields-A review.

4. Noby, N., Hussein, A., Saeed, H., and Embaby, A. M. (2020) “Recombinant cold-adapted halotolerant, organic solvent-stable esterase (estHIJ) from Bacillus halodurans. Analytical Biochemistry 591, 113554

5. Mangiagalli, M., Brocca, S., Orlando, M., and Lotti, M. (2020) The “cold revolution”. Present and future applications of cold-active enzymes and ice-binding proteins. N Biotechnol 55, 5–11

6. Gupta, S. K., Kataki, S., Chatterjee, S., Prasad, R. K., Datta, S., Vairale, M. G., Sharma, S., Dwivedi, S. K., and Gupta, D. K. (2020) Cold adaptation in bacteria with special focus on cellulase production and its potential application. Journal of Cleaner Production 258, 120351

7. Bornscheuer, U. T. (2002) Microbial carboxyl esterases: classification, properties and application in biocatalysis. FEMS Microbiol Rev 26, 73–81

8. Jaeger, K. E., Dijkstra, B. W., and Reetz, M. T. (1999) Bacterial biocatalysts: molecular biology, three-dimensional structures, and biotechnological applications of lipases. Annu Rev Microbiol 53, 315–351

9. Chandra, P., Enespa, Singh R., and Arora, P. K. (2020) Microbial lipases and their industrial applications: a comprehensive review. Microb Cell Fact 19, 169

10. Joseph, B., Ramteke, P. W., and Thomas, G. (2008) Cold active microbial lipases: some hot issues and recent developments. Biotechnology advances 26, 457–470

11. Kuddus, M. (2015) Cold-active microbial enzymes. Biochem Physiol 4, e132

12. Elend, C., Schmeisser, C., Hoebenreich, H., Steele, H., and Streit, W. (2007) Isolation and characterization of a metagenome-derived and cold-active lipase with high stereospecificity for (R)-ibuprofen esters. Journal of biotechnology 130, 370–377

13. Rotticci, D., Ottosson, J., Norin, T., and Hult, K. (2001) Candida antarctica Lipase BA Tool for the Preparation of Optically Active Alcohols. in Enzymes in Nonaqueous Solvents, Springer. pp 261–276

14. Maharana, A., and Ray, P. (2015) A novel cold-active lipase from psychrotolerant Pseudomonas sp. AKM-L5 showed organic solvent resistant and suitable for detergent formulation. Journal of Molecular Catalysis B: Enzymatic 120, 173–178

15. Santiago, M., Ramírez-Sarmiento, C. A., Zamora, R. A., and Parra, L. P. (2016) Discovery, molecular mechanisms, and industrial applications of cold-active enzymes. Frontiers in microbiology 7, 1408

16. Kuddus, M. (2018) Cold-active enzymes in food biotechnology: An updated mini review. J. Appl. Biol. Biotechnol 6, 58–63

17. Feller, G., and Gerday, C. (2003) Psychrophilic enzymes: hot topics in cold adaptation. Nat Rev Microbiol 1, 200–208

18. Gianese, G., Bossa, F., and Pascarella, S. (2002) Comparative structural analysis of psychrophilic and meso- and thermophilic enzymes. Proteins 47, 236–249

19. Siddiqui, K. S., and Cavicchioli, R. (2006) Cold-adapted enzymes. Annu Rev Biochem 75, 403–433

20. Feller, G. (2010) Protein stability and enzyme activity at extreme biological temperatures. J Phys Condens Matter 22, 323101

21. Marx, J., Collins, T., D’Amico, S., Feller, G., and Gerday, C. (2007) Cold-adapted enzymes from marine Antarctic microorganisms. Marine biotechnology 9, 293–304

22. Barroca, M., Santos, G., Gerday, C., and Collins, T. (2017) Biotechnological aspects of cold-active enzymes. in Psychrophiles: From Biodiversity to Biotechnology, Springer. pp 461–475

23. Zamora, R. A., Ramirez-Sarmiento, C. A., Castro-Fernandez, V., Villalobos, P., Maturana, P., Herrera-Morande, A., Komives, E. A., and Guixe, V. (2020) Tuning of Conformational Dynamics Through Evolution-Based Design Modulates the Catalytic Adaptability of an Extremophilic Kinase. ACS Catalysis 10, 10847–10857

24. Smalas, A. O., Leiros, H. K., Os, V., and Willassen, N. P. (2000) Cold adapted enzymes. Biotechnol Annu Rev 6, 1–57

25. Tutino, M. L., di Prisco, G., Marino, G., and de Pascale, D. (2009) Cold-adapted esterases and lipases: from fundamentals to application. Protein Pept Lett 16, 1172–1180

26. Kumar, S., and Nussinov, R. (2004) Different roles of electrostatics in heat and in cold: adaptation by citrate synthase. Chembiochem 5, 280–290

27. Mangiagalli, M., and Lotti, M. (2021) Cold-Active β-Galactosidases: Insight into Cold Adaption Mechanisms and Biotechnological Exploitation. Marine Drugs 19, 43

28. Mangiagalli, M., Lapi, M., Maione, S., Orlando, M., Brocca, S., Pesce, A., Barbiroli, A., Camilloni, C., Pucciarelli, S., and Lotti, M. (2021) The co-existence of cold activity and thermal stability in an Antarctic GH42 β-galactosidase relies on its hexameric quaternary arrangement. The FEBS Journal 288, 546–565

29. Noby, N., Saeed, H., Embaby, A. M., Pavlidis, I. V., and Hussein, A. (2018) Cloning, expression and characterization of cold active esterase (EstN7) from Bacillus cohnii strain N1: A novel member of family IV. International journal of biological macromolecules 120, 1247–1255

30. Zhu, X., Larsen, N. A., Basran, A., Bruce, N. C., and Wilson, I. A. (2003) Observation of an arsenic adduct in an acetyl esterase crystal structure. J Biol Chem 278, 2008–2014

31. Arpigny, J. L., and Jaeger, K. E. (1999) Bacterial lipolytic enzymes: classification and properties. Biochem J 343 Pt 1, 177–183

32. Krissinel, E., and Henrick, K. (2007) Inference of macromolecular assemblies from crystalline state. J Mol Biol 372, 774–797

33. McKary, M. G., Abendroth, J., Edwards, T. E., and Johnson, R. J. (2016) Structural Basis for the Strict Substrate Selectivity of the Mycobacterial Hydrolase LipW. Biochemistry 55, 7099–7111

34. Palm, G. J., Fernandez-Alvaro, E., Bogdanovic, X., Bartsch, S., Sczodrok, J., Singh, R. K., Bottcher, D., Atomi, H., Bornscheuer, U. T., and Hinrichs, W. (2011) The crystal structure of an esterase from the hyperthermophilic microorganism Pyrobaculum calidifontis VA1 explains its enantioselectivity. Appl Microbiol Biotechnol 91, 1061–1072

35. De Santi, C., Leiros, H. K., Di Scala, A., de Pascale, D., Altermark, B., and Willassen, N. P. (2016) Biochemical characterization and structural analysis of a new cold-active and salt-tolerant esterase from the marine bacterium Thalassospira sp. Extremophiles 20, 323–336

36. Costantini, S., Colonna, G., and Facchiano, A. M. (2008) ESBRI: a web server for evaluating salt bridges in proteins. Bioinformation 3, 137–138

37. Jacobs, D. J., Rader, A. J., Kuhn, L. A., and Thorpe, M. F. (2001) Protein flexibility predictions using graph theory. Proteins 44, 150–165

38. McManus, T. J., Wells, S. A., and Walker, A. B. (2019) Salt bridge impact on global rigidity and thermostability in thermophilic citrate synthase. Phys Biol 17, 016002

39. Tian, W., Chen, C., Lei, X., Zhao, J., and Liang, J. (2018) CASTp 3.0: computed atlas of surface topography of proteins. Nucleic Acids Res 46, W363–W367

40. Jurrus, E., Engel, D., Star, K., Monson, K., Brandi, J., Felberg, L. E., Brookes, D. H., Wilson, L., Chen, J., Liles, K., Chun, M., Li, P., Gohara, D. W., Dolinsky, T., Konecny, R., Koes, D. R., Nielsen, J. E., Head-Gordon, T., Geng, W., Krasny, R., Wei, G. W., Holst, M. J., McCammon, J. A., and Baker, N. A. (2018) Improvements to the APBS biomolecular solvation software suite. Protein Sci 27, 112–128

41. Li, P. Y., Ji, P., Li, C. Y., Zhang, Y., Wang, G. L., Zhang, X. Y., Xie, B. B., Qin, Q. L., Chen, X. L., Zhou, B. C., and Zhang, Y. Z. (2014) Structural basis for dimerization and catalysis of a novel esterase from the GTSAG motif subfamily of the bacterial hormone-sensitive lipase family. J Biol Chem 289, 19031–19041

42. Levisson, M., Han, G. W., Deller, M. C., Xu, Q., Biely, P., Hendriks, S., Ten Eyck, L. F., Flensburg, C., Roversi, P., Miller, M. D., McMullan, D., von Delft, F., Kreusch, A., Deacon, A. M., van der Oost, J., Lesley, S. A., Elsliger, M. A., Kengen, S. W., and Wilson, I. A. (2012) Functional and structural characterization of a thermostable acetyl esterase from Thermotoga maritima. Proteins 80, 1545–1559

43. Rader, A. J., Hespenheide, B. M., Kuhn, L. A., and Thorpe, M. F. (2002) Protein unfolding: rigidity lost. Proc Natl Acad Sci U S A 99, 3540–3545

44. Wells, S. A., Crennell, S. J., and Danson, M. J. (2014) Structures of mesophilic and extremophilic citrate synthases reveal rigidity and flexibility for function. Proteins 82, 2657–2670

45. Radestock, S., and Gohlke, H. (2011) Protein rigidity and thermophilic adaptation. Proteins 79, 1089–1108

46. Jaenicke, R., and Bohm, G. (1998) The stability of proteins in extreme environments. Curr Opin Struct Biol 8, 738–748

47. Jimenez-Roldan, J. E., Freedman, R. B., Romer, R. A., and Wells, S. A. (2012) Rapid simulation of protein motion: merging flexibility, rigidity and normal mode analyses. Phys Biol 9, 016008

48. Wells, S., Menor, S., Hespenheide, B., and Thorpe, M. F. (2005) Constrained geometric simulation of diffusive motion in proteins. Phys Biol 2, S127–136

49. Fu, J., Leiros, H. K., de Pascale, D., Johnson, K. A., Blencke, H. M., and Landfald, B. (2013) Functional and structural studies of a novel cold-adapted esterase from an Arctic intertidal metagenomic library. Appl Microbiol Biotechnol 97, 3965–3978

50. Miyazaki, K., Wintrode, P. L., Grayling, R. A., Rubingh, D. N., and Arnold, F. H. (2000) Directed evolution study of temperature adaptation in a psychrophilic enzyme. J Mol Biol 297, 1015–1026

51. Merlino, A., Russo Krauss, I., Castellano, I., De Vendittis, E., Rossi, B., Conte, M., Vergara, A., and Sica, F. (2010) Structure and flexibility in cold-adapted iron superoxide dismutases: the case of the enzyme isolated from Pseudoalteromonas haloplanktis. J Struct Biol 172, 343–352

52. Pischedda, A., Ramasamy, K. P., Mangiagalli, M., Chiappori, F., Milanesi, L., Miceli, C., Pucciarelli, S., and Lotti, M. (2018) Antarctic marine ciliates under stress: superoxide dismutases from the psychrophilic Euplotes focardii are cold-active yet heat tolerant enzymes. Sci Rep 8, 14721

53. Karan, R., Mathew, S., Muhammad, R., Bautista, D. B., Vogler, M., Eppinger, J., Oliva, R., Cavallo, L., Arold, S. T., and Rueping, M. (2020) Understanding High-Salt and Cold Adaptation of a Polyextremophilic Enzyme. Microorganisms 8

54. Arcus, V. L., van der Kamp, M. W., Pudney, C. R., and Mulholland, A. J. (2020) Enzyme evolution and the temperature dependence of enzyme catalysis. Curr Opin Struct Biol 65, 96–101

55. Smith, D. K., Radivojac, P., Obradovic, Z., Dunker, A. K., and Zhu, G. (2003) Improved amino acid flexibility parameters. Protein Sci 12, 1060–1072

56. Lonhienne, T., Gerday, C., and Feller, G. (2000) Psychrophilic enzymes: revisiting the thermodynamic parameters of activation may explain local flexibility. Biochim Biophys Acta 1543, 1–10

57. Kim, S. Y., Hwang, K. Y., Kim, S. H., Sung, H. C., Han, Y. S., and Cho, Y. (1999) Structural basis for cold adaptation. Sequence, biochemical properties, and crystal structure of malate dehydrogenase from a psychrophile Aquaspirillium arcticum. J Biol Chem 274, 11761–11767

58. Chovancova, E., Pavelka, A., Benes, P., Strnad, O., Brezovsky, J., Kozlikova, B., Gora, A., Sustr, V., Klvana, M., Medek, P., Biedermannova, L., Sochor, J., and Damborsky, J. (2012) CAVER 3.0: a tool for the analysis of transport pathways in dynamic protein structures. PLoS Comput Biol 8, e1002708

59. Paredes, D. I., Watters, K., Pitman, D. J., Bystroff, C., and Dordick, J. S. (2011) Comparative void-volume analysis of psychrophilic and mesophilic enzymes: structural bioinformatics of psychrophilic enzymes reveals sources of core flexibility. BMC structural biology 11, 1–9

60. McCoy, A. J., Grosse-Kunstleve, R. W., Adams, P. D., Winn, M. D., Storoni, L. C., and Read, R. J. (2007) Phaser crystallographic software. J Appl Crystallogr 40, 658–674

61. Murshudov, G. N., Vagin, A. A., and Dodson, E. J. (1997) Refinement of macromolecular structures by the maximum-likelihood method. Acta Crystallogr D Biol Crystallogr 53, 240–255

62. Murshudov, G. N., Skubak, P., Lebedev, A. A., Pannu, N. S., Steiner, R. A., Nicholls, R. A., Winn, M. D., Long, F., and Vagin, A. A. (2011) REFMAC5 for the refinement of macromolecular crystal structures. Acta Crystallogr D Biol Crystallogr 67, 355–367

63. Emsley, P., and Cowtan, K. (2004) Coot: model-building tools for molecular graphics. Acta Crystallogr D Biol Crystallogr 60, 2126–2132

64. Suhre, K., and Sanejouand, Y. H. (2004) ElNemo: a normal mode web server for protein movement analysis and the generation of templates for molecular replacement. Nucleic Acids Res 32, W610–614

65. Pronk, S., Pall, S., Schulz, R., Larsson, P., Bjelkmar, P., Apostolov, R., Shirts, M. R., Smith, J. C., Kasson, P. M., van der Spoel, D., Hess, B., and Lindahl, E. (2013) GROMACS 4.5: a high-throughput and highly parallel open source molecular simulation toolkit. Bioinformatics 29, 845–854

66. Lindorff-Larsen, K., Piana, S., Palmo, K., Maragakis, P., Klepeis, J. L., Dror, R. O., and Shaw, D. E. (2010) Improved side-chain torsion potentials for the Amber ff99SB protein force field. Proteins 78, 1950–1958

