## Supporting Table and figures for "Structure and dynamics of a cold-active esterase reveals water entropy and active site accessibility as the likely drivers for cold-adaptation"

**Supporting Information.**

**Supporting Table S1.** Data collection and refinement statistics for EstN7

| **PDB Entry** | |
|---|---|
| **Data Collection** | |
| Diamond Beamline | I03 |
| Date | 2019-04-18 |
| Wavelength | 0.97629 |
| **Crystal Data** | |
| Crystallisation Conditions | 0.2M lithium chloride, 0.1M sodium |
| *a,b,c* (Å) | 109.91, 109.91, 126.76 |
| $\alpha=\beta=\gamma$ (°) | 90.0, 90.0, 90.0 |
| Space group | $P\ 4_3\ 2_1\ 2$ |
| Resolution (Å) | 1.61 – 83.04 |
| Outer shell | 1.61 – 1.65 |
| *R*-merge (%) | 10.6 (219.1) |
| *R-pim (%)* | 2.9 (63.4) |
| *R*-meas (%) | 11.0 (234.9) |
| CC1/2 | 0.999 (0.672) |
| I / σ(I) | 13.8 (1.0) |
| Completeness (%) | 100.0 (100.0) |
| Multiplicity | 14.5 (13.7) |
| Total Measurements | 1,459,945 (100,341) |
| Unique Reflections | 100,631 (7,339) |
| Wilson B-factor(Å$^2$) | 21.4 |
| **Refinement Statistics** | |
| Total number of refined atoms | 5591 |
| R-work reflections | 95,500 |
| R-free reflections | 5,029 |
| R-work/R-free (%) | 14.4 / 18.7 |
| **rms deviations** | |
| Bond lengths (Å) | 0.013 |
| Bond Angles (°) | 1.739 |
| [b]Coordinate error | 0.056 |
| Mean B value (Å$^2$) | 30.4 |
| **Ramachandran Statistics** | |
| Favoured/allowed/Outliers | 565 / 20 / 3 |
| % | 96.1 / 3.4 / 0.5 |

[a] Figures in brackets refer to outer resolution shell, where applicable.
[b] Coordinate Estimated Standard Uncertainty in (Å), calculated based on maximum likelihood statistics.

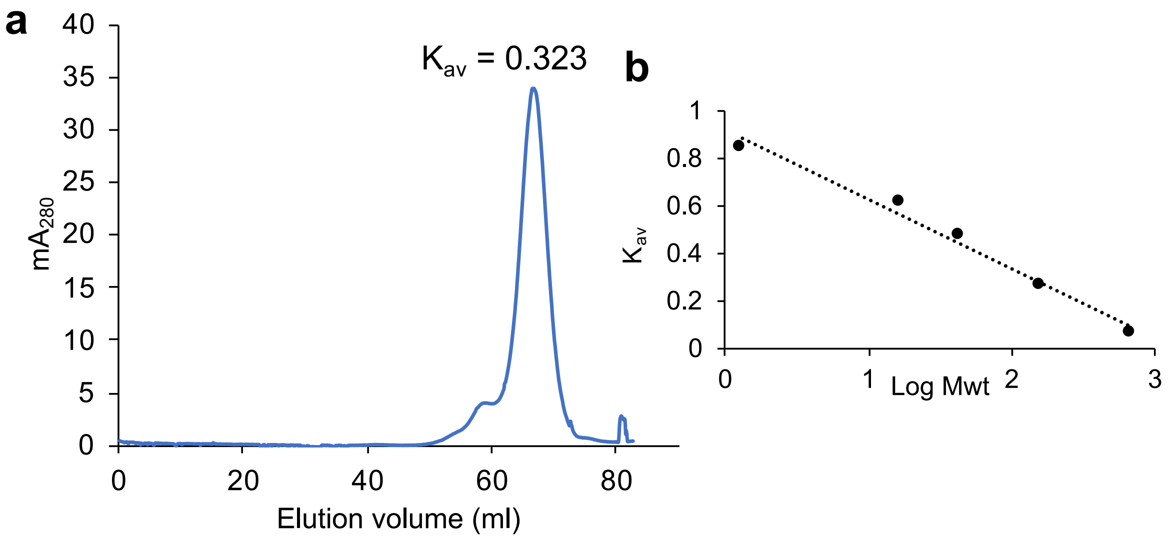


**Supporting Figure S1.** Quaternary structure analysis by size exclusion chromatography. (a) Elution profile of EstN7 from a Hiload™ 16/60 Superdex™ S200 pg column (Cytiva). The calculated $K_{av}$ was 0.323 based on the equation $K_{av} = (V_e-V_0) / (V_t-V_0)$ where $V_e$ is the peak elution volume, $V_0$ is void volume (42 ml) and $V_t$ is total column volume (120 ml). Absorbance was measured at 280 nm. (b) Calibration curve for the column with protein standards of molecular weight 670, 158, 44, 17 and 1.35 kDa. The estimated molecular mass for EstN7 is ~100 kDa, with the monomeric mass being 37kDa.

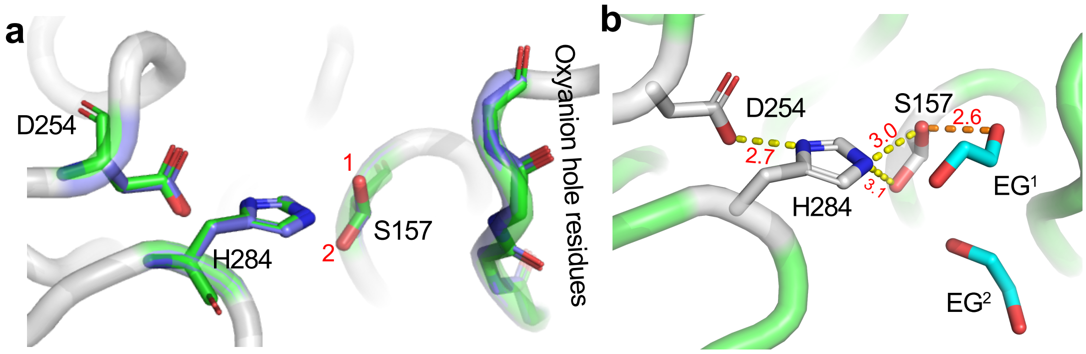


**Supporting Figure S2.** Catalytic triad and oxyanion hole arrangement. (a) Overlay of the catalytic triad oxyanion hole from subunit A (blue) and subunit B (green), with the two rotamers of S157 labelled as 1 and 2. (b) arrangement of ethylene glycol

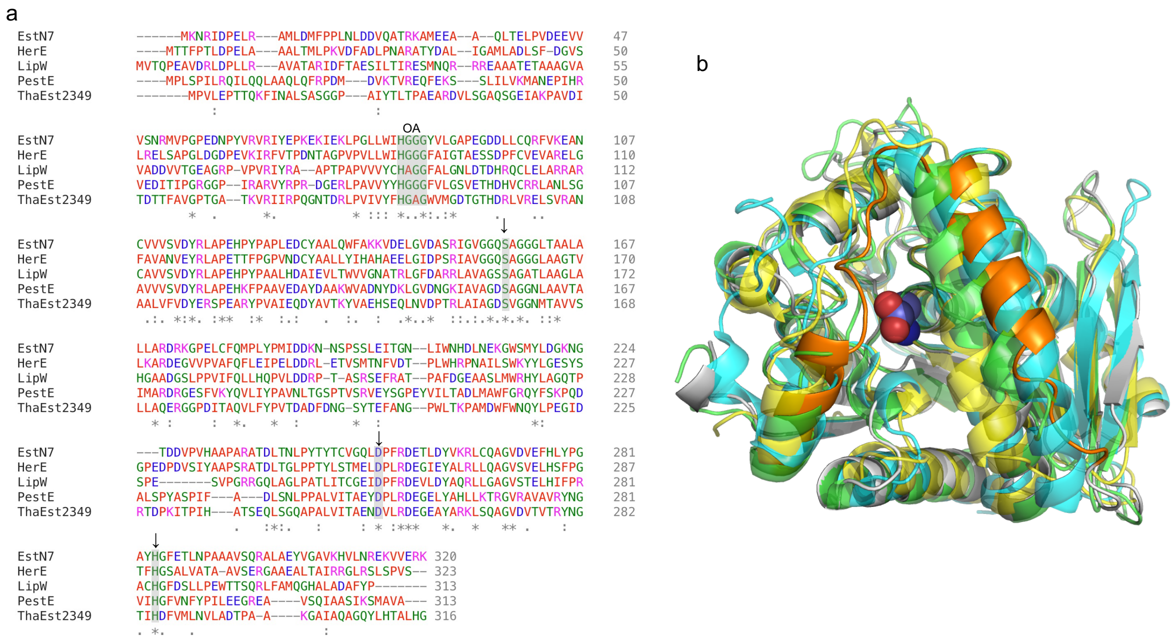


**Figure S3.** Sequence and structural alignment of close structural homologues. (a) sequence alignment of EstN7, HerE (heroin esterase) from *Rhodococcus* sp. strain H1 (1), a mycobacterial LipW (2), PestE from the thermophile *Pyrobaculum calidifontis* (3) and a putative cold-adapted esterase from marine bacterium Thalassospira sp (4). Catalytic residues are indicated by the arrows and the oxyanion hole residues by the grey box annotated with OA. Acidic residues are coloured blue, basic residue pink, polar residues green and hydrophobic red. The "*", ":" and "." refer to conserved, strong conserved and weakly conserved residues, respectively. (b) Structural alignment of EstN7 (grey with cap region coloured orange and the serine nucleophile S157 shown as purple spheres), HerE (PDB 1lzk; coloured green), LipW (PDB 3qh4; coloured cyan) and PestE (PDB 2yh2; coloured yellow).

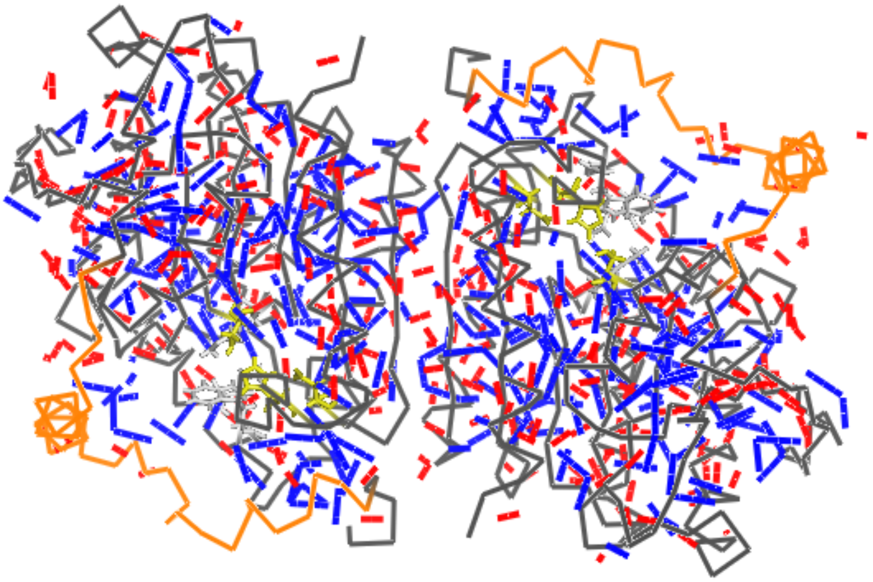

**Figure S4.** Non-covalent constraints in the crystal structure of EstN7. The protein backbone is shown as grey ribbon; the cap region is highlighted in orange. The catalytic triad residues are shown as yellow sticks and the plug residues as white sticks. Blue bars indicate hydrophobic tether interactions, in which two nonpolar nonhydrogen atoms lie at a distance less than 0.5Å greater than the sum of their steric radii. Red bars indicate polar interactions (donor-hydrogen-acceptor) with an effective strength of -4.0 kcal/mol or better according to their geometry(5,6).

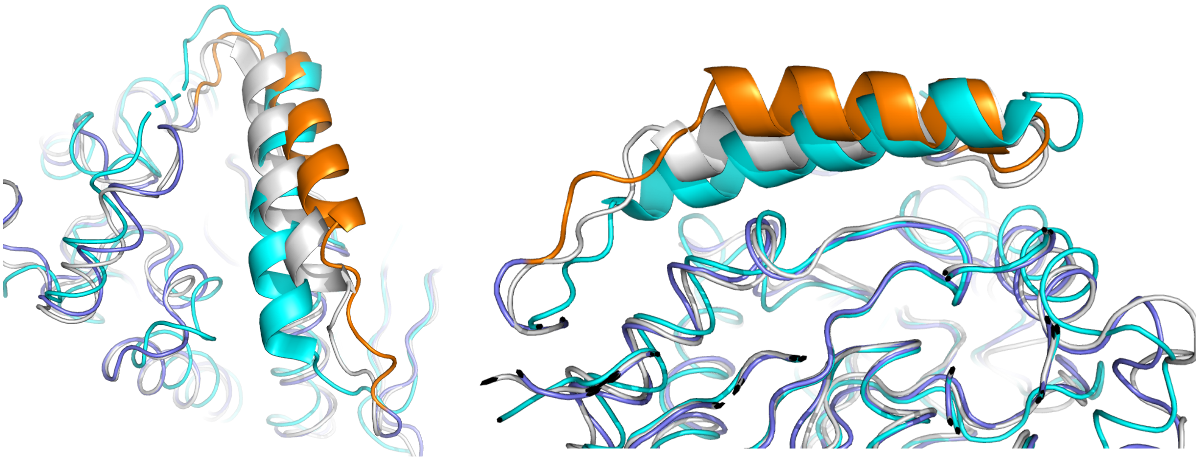

**Figure S5** Comparison of the second helical component of the cap structure from EstN7 (orange and blue), HerE (1) (grey; PDB 1LZK) and PestE (3) (cyan; PDB 2YH2).

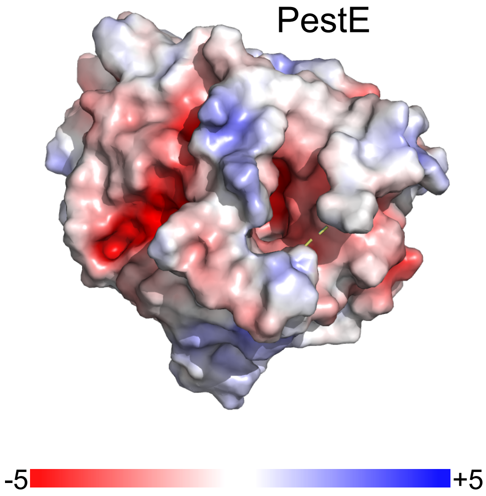


**Figure S6.** Surface electrostatic profile of thermophilic EstN7-related esterase PestE subunit A. Electrostatic potential surface of was calculated using APBS (7) with the protein orientation identical to that in Figure 2b. Colour scaling of electrostatic potential is shown.

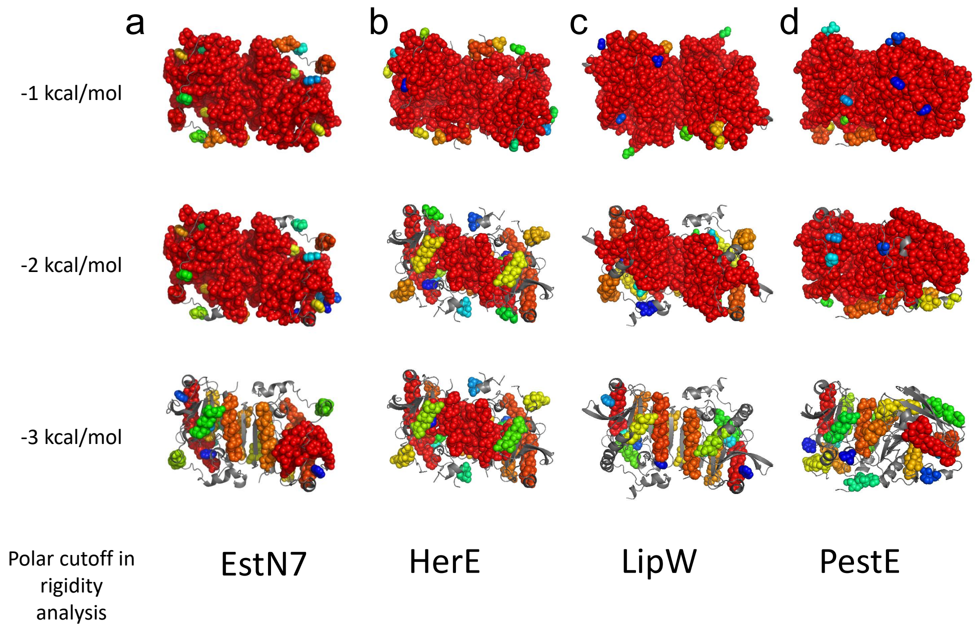


**Figure S7.** Comparative rigidity analysis of (a) EstN7 with (b) HerE, (c) LipW and (d) PestE. The 20 largest rigid clusters are shown as spheres and coloured as rainbow from red (largest) to blue (20$^{th}$ largest). Polar cutoff values of -1, -2 and -3 kcal/mol are shown. Flexible regions are shown as backbone cartoon and coloured grey. The analysis of EstN7 shown here uses 'ambient' settings, and the structure thus appears comparably rigid to PestE. See main manuscript Figure 2(c) for a 'cold' analysis allowing for the weakening of the hydrophobic effect at low temperatures.

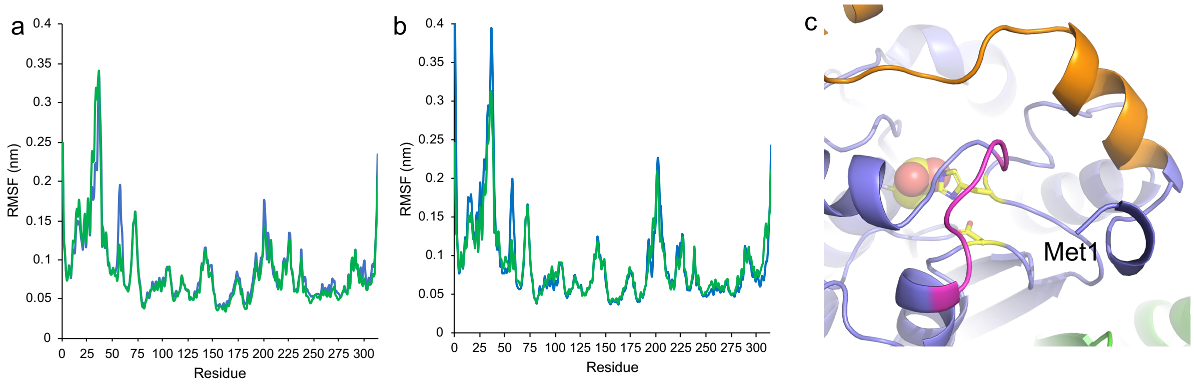


**Figure S8.** Molecular dynamics at different temperatures. RMSF over the A (blue) and B (green) subunit at (a) 283K (10°C) and (b) 308K (35°C). The simulations were run over 100 ns. The RMSF values are the average of 4 independent MD simulations. (c) Region E199-I205 that has temperature dependent dynamic profile (see main manuscript, Figure 3). The proximity to the N-terminal (Met1), is shown. The cap region is coloured orange and the catalytic triad yellow.

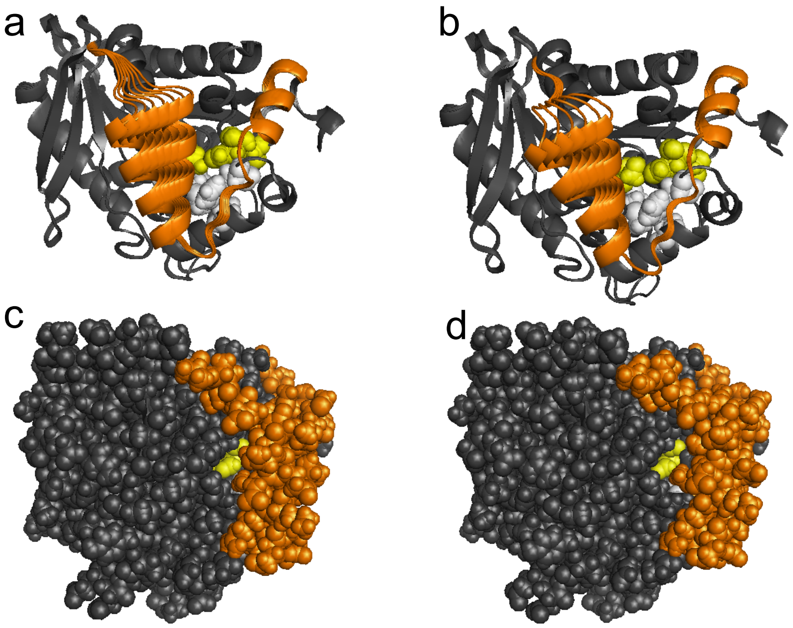


**Figure S9.** Visualisation of geometric simulations of flexible motion(8) of EstN7 along low-frequency normal mode directions localised principally on the cap regions: (a) view of the cap and active site of chain A, showing an overlay of frames from motion along normal mode 10; (b) view of the cap and active site of chain B, showing an overlay of frames from motion along normal mode 09; (c) all-atom view of chain A showing cap (orange) and active site (yellow) residues, rotated to show the suggested alternate access channel in the crystal structure; (d) all-atom view of chain A showing cap (orange) and active site (yellow) residues, as in (c), showing a conformation from flexible motion biased along normal mode 10. Motion of the flexible cap exposes the active site to view.

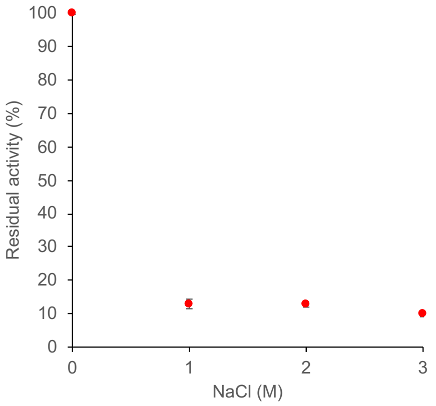


**Figure S10.** Halotolerance of EstN7. The activity of EstN7 was performed as described previously (9) in the presence of 0, 1, 2 and 3 M NaCl. All sample where incubated at room temperature for 30 min prior to activity measurements. The percentage residual activity is relative to the 0 M NaCl sample. Each measurement was performed in triplicate.

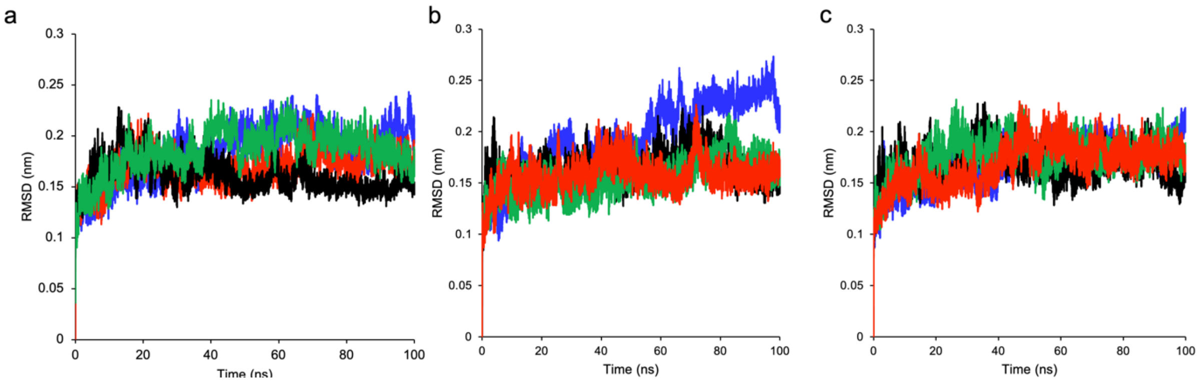


**Figure S11.** RMSD at (a) 298K, (b) 283K and (c) 308K over the course of 100 ns for each simulation. There are 4 separate simulations coloured black, green, blue and red that comprise each MD analysis.

### Supporting References


1. Zhu, X., Larsen, N. A., Basran, A., Bruce, N. C., and Wilson, I. A. (2003) Observation of an arsenic adduct in an acetyl esterase crystal structure. *J Biol Chem* **278**, 2008-2014
2. McKary, M. G., Abendroth, J., Edwards, T. E., and Johnson, R. J. (2016) Structural Basis for the Strict Substrate Selectivity of the Mycobacterial Hydrolase LipW. *Biochemistry* **55**, 7099-7111
3. Palm, G. J., Fernandez-Alvaro, E., Bogdanovic, X., Bartsch, S., Sczodrok, J., Singh, R. K., Bottcher, D., Atomi, H., Bornscheuer, U. T., and Hinrichs, W. (2011) The crystal structure of an esterase from the hyperthermophilic microorganism Pyrobaculum calidifontis VA1 explains its enantioselectivity. *Appl Microbiol Biotechnol* **91**, 1061-1072
4. De Santi, C., Leiros, H. K., Di Scala, A., de Pascale, D., Altermark, B., and Willassen, N. P. (2016) Biochemical characterization and structural analysis of a new cold-active and salt-tolerant esterase from the marine bacterium Thalassospira sp. *Extremophiles* **20**, 323-336
5. McManus, T. J., Wells, S. A., and Walker, A. B. (2019) Salt bridge impact on global rigidity and thermostability in thermophilic citrate synthase. *Phys Biol* **17**, 016002
6. Jacobs, D. J., Rader, A. J., Kuhn, L. A., and Thorpe, M. F. (2001) Protein flexibility predictions using graph theory. *Proteins* **44**, 150-165
7. Jurrus, E., Engel, D., Star, K., Monson, K., Brandi, J., Felberg, L. E., Brookes, D. H., Wilson, L., Chen, J., Liles, K., Chun, M., Li, P., Gohara, D. W., Dolinsky, T., Konecny, R., Koes, D. R., Nielsen, J. E., Head-Gordon, T., Geng, W., Krasny, R., Wei, G. W., Holst, M. J., McCammon, J. A., and Baker, N. A. (2018) Improvements to the APBS biomolecular solvation software suite. *Protein Sci* **27**, 112-128
8. Jimenez-Roldan, J. E., Freedman, R. B., Romer, R. A., and Wells, S. A. (2012) Rapid simulation of protein motion: merging flexibility, rigidity and normal mode analyses. *Phys Biol* **9**, 016008
9. Noby, N., Saeed, H., Embaby, A. M., Pavlidis, I. V., and Hussein, A. (2018) Cloning, expression and characterization of cold active esterase (EstN7) from Bacillus cohnii strain N1: A novel member of family IV. *International journal of biological macromolecules* **120**, 1247-1255
